# Developmental transcriptomic analysis of cultured primary mouse cortical neurons reveals sex-specific expression of neuropeptides

**DOI:** 10.64898/2026.04.21.719935

**Authors:** Alekh Paranjapye, Rili Ahmad, John J. Gerace, Erica Korb

**Author notes:** Corresponding author: Erica Korb.

## Abstract

Primary neuronal cultures derived from mouse tissue serve as an essential model for investigating neuronal development and function. Despite this, comprehensive developmental and sex-specific transcriptomic profiles in primary neurons have not been defined. Here, we performed multiplexed RNA-sequencing of neurons derived from male and female cortices across time. We validated this approach by assessing established neuronal maturation genes and we identified highly stable genes to serve as controls across development. Next, using linear modeling and temporal regression, we defined developmentally regulated transcripts, longitudinal expression dynamics, and gene signatures associated with transition states throughout development. Unexpectedly, this also revealed sex-specific effects on autosomal genes that emerge only after neuronal maturation even in the absence of *in vivo* cues. Most notably, neuropeptide genes Cortistatin and Neurokinin A are more highly expressed in female neurons. Furthermore, exposure to these neuropeptides elicited distinct transcriptional responses in male-versus female-derived cultures. These findings provide a valuable resource and reveal sex-specific autosomal transcriptional signatures that emerge in neurons maintained *ex vivo*.

**Highlights:**

- Multiplexed RNA-sequencing of primary cultured neurons across neuronal maturation provides a new resource for the field.
- Gene signatures associated with transition states are identified through linear modeling.
- Sex-specific regulation of autosomal genes encoding neuropeptides emerge even in the absence of *in vivo* cues.
- Exposure to neuropeptides elicit distinct transcriptional responses in male and female primary neurons.

## Introduction

Primary neuron cultures derived from embryonic mice are widely used *in vitro* models for studying developmental neurobiology in a controlled environment, independent of social, environmental, and other *in vivo* influences^1–13^. These systems overcome limitations associated with isolating neurons from heterogeneous brain tissue, which can result in the loss of dendrites and axons and lead to sample degradation. They also avoid complications and mixed cell populations that may arise during the differentiation of pluripotent stem cells. Although primary culture systems have inherent limitations and lack the complex circuitry of *in vivo* models, they remain invaluable for testing drug responses^14,15^, genetic perturbations associated with diseases^16^ and disorders^17–19^, altered physiology^20,21,19^, and more. Additionally, primary cultures can be more readily manipulated and generated at scales that are more difficult to achieve than other neuron-selective methodologies, making them well-suited for detailed mechanistic studies.

Atlases of bulk and single-nuclei transcriptomic data have become increasingly important in neurobiology^22^. However, despite the widespread use of primary neuron cultures, relatively few RNA-sequencing datasets profile wildtype, untreated cells, and, to our knowledge, none include neurons derived from both male and female animals. Existing datasets are typically limited to one or two time points or track the differentiation of murine embryonic stem cells. While highly informative, prior work providing comprehensive RNA-sequencing trajectories^23^ focused on stem cell-derived neurons rather than tissue-isolated primary neurons and was limited to male cells. A comprehensive gene-expression dataset from primary neurons that captures developmental trajectories and sex-specific effects would therefore be a highly valuable resource. It will provide a reference for genes or pathways of interest, inform ideal windows of culture maturation for specific experimental questions, and help delineate the strengths, weaknesses, and *in vivo* relevance of primary neuronal models. Further, given that sex-specific effects are readily detectable in the brain^24–32^, such approaches have the potential to determine whether sex-specific differences are also detectable in the absence of *in vivo* influences.

To this end, we generated the first longitudinal, bulk RNA-sequencing data set for embryonic mouse neuron cultures. We profiled the transcriptome of ten embryos, five of each sex, at six time points from pre-culture through late maturation. Using one vs all, stepwise, and temporal regression analyses, we defined gene expression signatures that shift across time in culture, including sex-specific changes. These included novel, sex-dependent responses to neuropeptides. Finally, we provide the field with an openly accessible website (https://www.korblab.com/data) for viewing and downloading data. Together, these findings highlight the importance of including sex as a variable, even in highly controlled culture systems, and provide a resource that defines gene expression across neuronal development.

## Results

### Transcriptome of primary cultured mouse cortical neurons throughout maturation

We sought to fill an outstanding gap in the field by defining the transcriptome of primary cultured cortical neurons across neuronal maturation and account for sex-specific effects. We cultured neurons derived from 5 female and 5 male embryonic day 16.5 mouse cortices. RNA was isolated at 6 time points: immediately after tissue dissociation and prior to plating (day 0), and after plating at 1, 5, 10, 15, and 20 days in vitro (DIVs). We selected DIV20 as the final time point because neuronal health becomes inconsistent beyond three weeks with standard protocols, and based on standard time points found in the literature^4,7,9,10^. We then processed all samples for multiplexed RNA-sequencing using Bulk RNA barcoding and sequencing (BRB-seq)^33^ (Fig. 1A), which indexes samples prior to sequencing library preparation, allowing for immediate pooling to reduce variability, minimize cost, and increase throughput.

**Figure 1:**
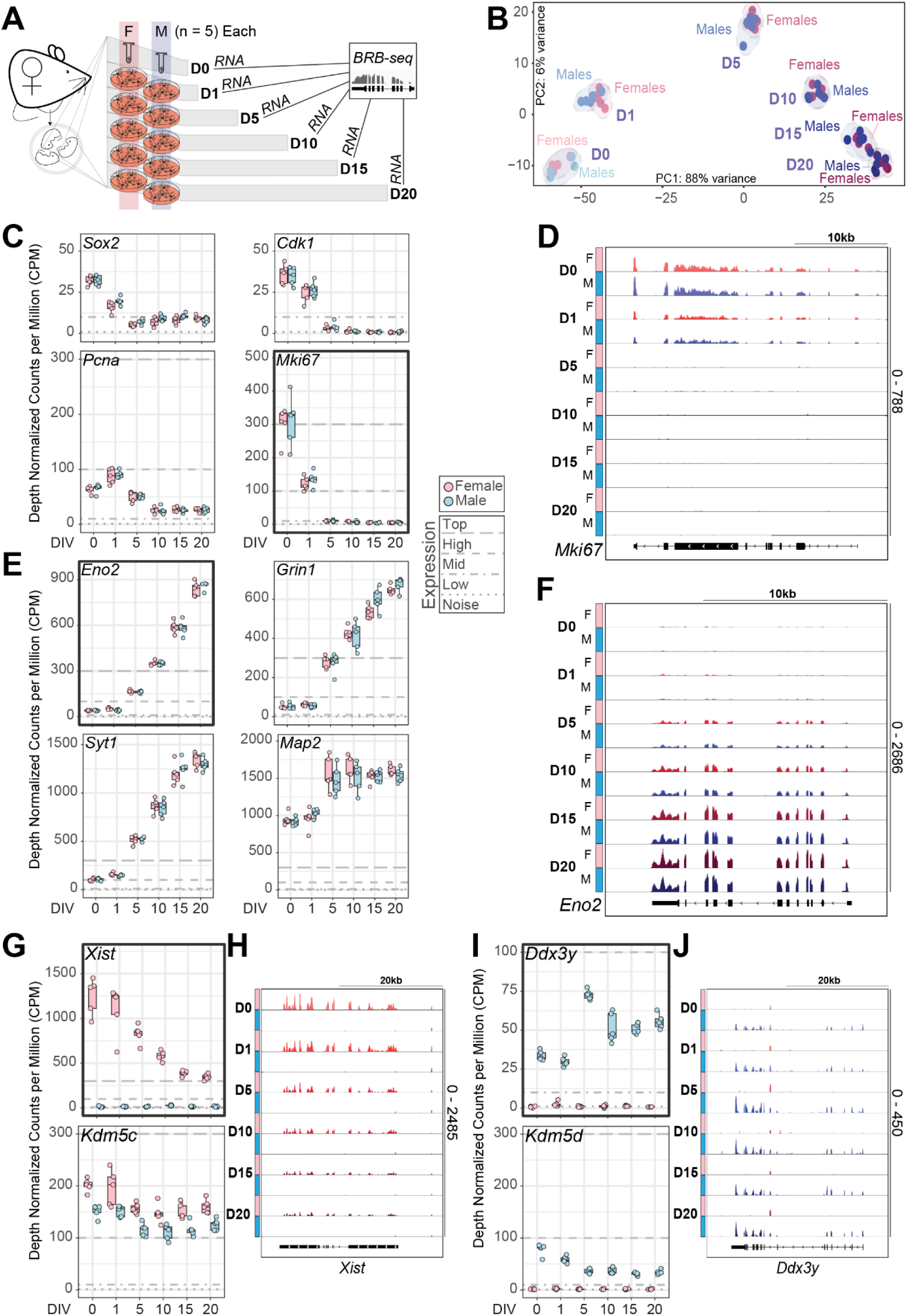
RNA-sequencing of a primary embryonic mouse cortical culture time course. A. Schematic of the experimental timeline for the comparison of transcriptomes of female- and male-derived neurons at six time points. B. Principal component analysis of the first two PCs with each replicate colored by sex. C. Counts per million (CPM) box plots of the expression of genes downregulated over days *in vitro* (n = 5 per sex). Bolded lines demarcate thresholds of very high to high (CPM > 300), high to mid (300 > CPM > 100), mid to low (100 > CPM > 10), and low to noise limit expression (10 > CPM). D. Gene track of aligned transcripts over the *Mki67* gene. Replicates by time and sex are overlayed for each track. E. CPM box plots of the expression of genes upregulated over days *in vitro*. F. Gene track of aligned transcripts over the *Eno2* gene. G. CPM box plots of the expression of female-specific genes. H. Gene track of aligned transcripts over the *Xist* gene. I. CPM box plots of the expression of genes on the Y-chromosome. J. Gene track of aligned transcripts over the *Ddx3y*.

We detected no outliers across replicates and observed an expected developmental trajectory based on principal component analysis (PCA) (Fig. 1B). To verify that the cultured neurons exhibited gene expression changes consistent with neuronal maturation, we assessed expression of four genes known to decrease during neuronal development and maturation (*Sox2*, *Cdk1*, *Pcna*, *Mki67*)^34–37^ and four known to increase (*Eno2*, *Grin1*, *Syt1*, *Map2*)^38–41^ (Fig 1C-F). The first group decreased in expression as expected over time, with *Cdk1* and *Mki67* falling below the noise threshold. In contrast, *Eno2*, *Grin1*, and *Syt1* increased steadily over time. *Map2* rapidly increased between D1 and D5 to reach high expression. We further confirmed that BRB-seq exhibits full exonic coverage and limited 3’ bias (Fig 1D, 1F). To validate PCR-based genotyping for pup sex, we confirmed anticipated expression patterns of two X-linked genes that have greater expression in females, *Xist* and *Kdm5c* (Fig. 1G-H). Similarly, we confirmed that expression of genes found on the Y chromosome, such as *Ddx3y* and *Kdm5d* are only detectable in the male replicates (Fig. 1I-J).

Next, we evaluated which cortical cell types were represented in neuronal cultures over time. Using a young adult mouse single-nuclei RNA-seq cortex dataset, we estimated the proportions of our bulk RNA-seq attributable to each of the mature neuronal and non-neuronal cell types (Supplemental Fig. 1A). Imputation was performed using the average single-nuclei gene expression across each of six excitatory neuron populations, six inhibitory neuron populations, and four non-neuronal cell populations against the normalized gene expression of each bulk sample at each time point. We found that cultures most closely matched the expression profiles of cortical excitatory neurons, as well as Pvalb+ and one subset of glycine/GABA inhibitory neurons, with greatest statistical confidence at DIV15 and DIV20. As expected, the proportion of the bulk dataset that could be explained by glia or radial glial populations decreased after plating (beyond DIV 0) and became undetectable after AraC treatment (performed at DIV 3) which removes dividing cells.

Having validated that neuronal maturation genes change as anticipated, X and Y genes are expressed in female and male neurons, and that expected cell types are present in primary cultures, we sought to make this resource readily available for widespread use. We generated a website that allows for any genes of interest to be visualized with downloadable graphs. Further, count data from either selected genes or all genes can similarly be downloaded for subsequent plotting independently. This resource is are available at https://www.korblab.com/data.

### Time-point-specific gene expression profiles

Selecting an appropriate time point for analysis is essential and requires an understanding of the unique expression profiles associated with each maturation stage in neuronal culture experiments. We therefore sought to identify differentially expressed genes (DEGs) at each time point relative to all other time points using EdgeR and limma (Fig. 2A-C). As expected, the greatest number of DEGs was observed at D0 (compared to all post-plating time points), when cells have recently been exposed to the *in vivo* environment, tissue has just undergone dissection and dissociation, and culturing conditions and media components have yet to enrich for specific cell populations. Gene ontology analysis of DEGs revealed substantial enrichment for transcripts encoding proteins involved in chromatin remodeling and the nuclear speck, with numerous writers and erasers of histone modifications, such as *Smarcc2*, *Dot1l*, *Ehmt2*, *Kmt2e*, *Setd2*, and *Setd5*, showing highest expression at D0 compared to later stages. In contrast, mitochondrial components, particularly those involved in translation and transport (Supplementary Fig. 2A, Supplementary Table 1-2), were expressed at lower levels. Among those genes upregulated at D1, corresponding to the transition to culture and the earliest phases of neuronal maturation, we observed significant increases in transcripts encoding ribosomal and other translation-associated proteins.

**Figure 2:**
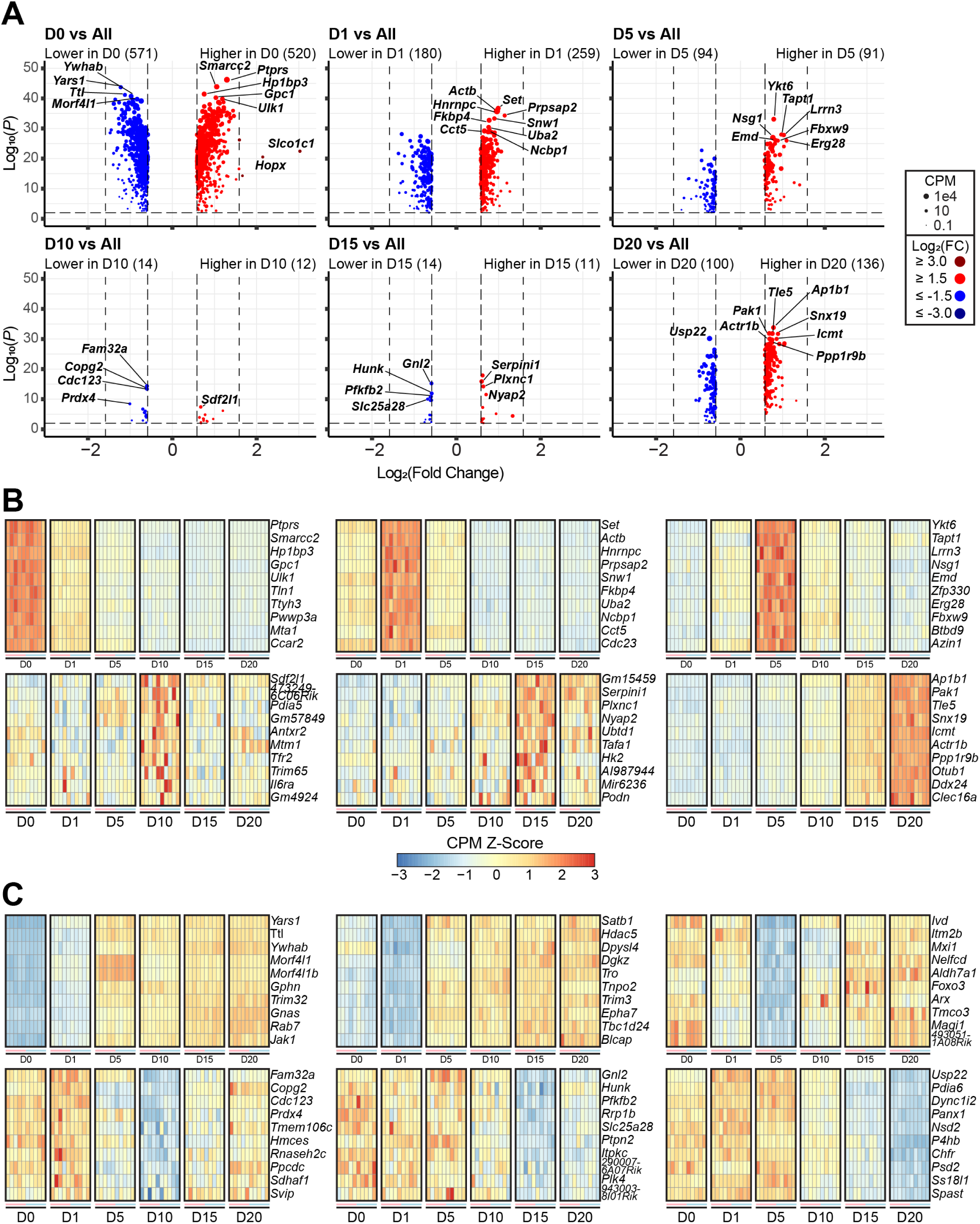
Timepoint-specific differential gene expression analysis of primary cultures. A. Volcano plots of differentially expressed genes (DEGs) comparing each time point against the regression aggregate of all other time points. Points are scaled in size proportional to its CPM and colored by fold change. B-C. Heatmaps of the top 10 upregulated (B) or downregulated (C) DEGs in the timepoint-specific comparisons.

Decreasing numbers of time-point-specific transcripts were identified across subsequent days *in vitro*, with the fewest observed at D10 and D15. However, transcripts specifically upregulated at D1 were enriched for genes involved in RNA processing (Supplemental Fig. 2B). Interestingly, although few significant DEGs were unique to the middle time points, we unexpectedly detected a substantial number at DIV20 (Fig. 2A). No significant gene ontology groups were enriched among the D20-specific genes, but several transcripts encoding components of the trans-golgi network (*Ap1b1*, *Ap5s1*, *M6pr*, *Gcc2*, *Cog4*) were upregulated. This network is essential in the establishment, organization, and functioning of neuronal circuitry^42,43^. We also observed increased expression of numerous genes involved in synaptic signaling, including *Ap1b1*, a member of the clathrin associated adaptor protein 1 (AP-1) complex, *Pak1*, a serine/threonine kinase, and *Mtor,* which is involved in brain circuitry and extracellular signaling responses^44^. These findings suggest that the re-emergence of transcriptomic differences at later time points may reflect heightened neuronal activity at more advanced stages *in vitro*^19^.

We next sought to identify genes that exhibit minimal expression changes throughout neuronal development, such as those commonly used for normalization across samples in qRT-PCR analysis. We first examined commonly used housekeeping genes including *Actb* (actin), *B2m* (beta-2-microglobulin), *Gapdh*, *Hprt1*, *Lama1 and Lmnb1* (Laminin), *Tubb2b* (beta-tubulin), and the 60S ribosomal component *Rplp1* (Supplemental Figure 3). Unexpectedly, every housekeeping gene exhibited marked gene expression changes across the time course, with the exception of *B2m*. Thus, while these genes may be valuable controls to compare experimental conditions *within* a time point, they are less likely to be useful when comparing across neuronal development. However, we identified other genes with stable and high expression across the time course by comparing the ratio between normalized expression and standard deviation across time. The topmost stable genes include *Dync1h1, Septin3, Trim2*, and *Copa*. *Dync1h1* and *Septin3* encode protein components of essential biological pathways (the dynein complex and cytoskeleton, respectively).^45,46^ *Trim2* encodes a ubiquitous ubiquitin ligase^47^ and *Copa* encodes a protein essential for intracellular trafficking^48^. These genes may provide more stable normalization factors than many standard housekeeping genes for assays testing transcript regulation across neuronal maturation.

### Defining differentially expressed genes at developmental transitions

Next, we performed sequential pairwise comparisons between each time point and the subsequent time point to identify the periods of greatest transcriptomic change and the genes involved (Fig. 3A). We detected up and downregulated DEGs in each comparison, with the greatest number detected at earlier stages of maturation. The transition from D0 to D1 was marked by upregulation of mitochondrial-associated transcripts and as much as an 8-fold downregulation of genes primarily involved in cell motility including the laminins (*Lama1*, *Lama2*, *Lama3*, *Lama4*, *Lama5*, *Lamb2*, *Lamc1*) and integrin subunits (*Itga3*, *Itga4*, *Itga5*, *Itga6*). Increased oxidative phosphorylation and mitochondrial function are hallmarks of neuronal development^49,50^ due to the high energy demands of the cell. The reduction in cytoskeleton components associated with motility may reflect the transition from migratory immature neurons present in the E16.5 brain to maturing and stationary neurons in culture (Supplemental Fig. 1). The D1 to D5 transition was similarly marked by significant transcriptional changes with nearly 5000 DEGs, with 181 showing over an 8-fold difference. Many of these genes were further modulated later in the time course, particularly at earlier maturation stages (Supplemental Fig. 4A). We also found genes such as potassium channel *Kcna2* and the GABA receptor *Gabra1* consistently upregulated across all five transitions.

**Figure 3:**
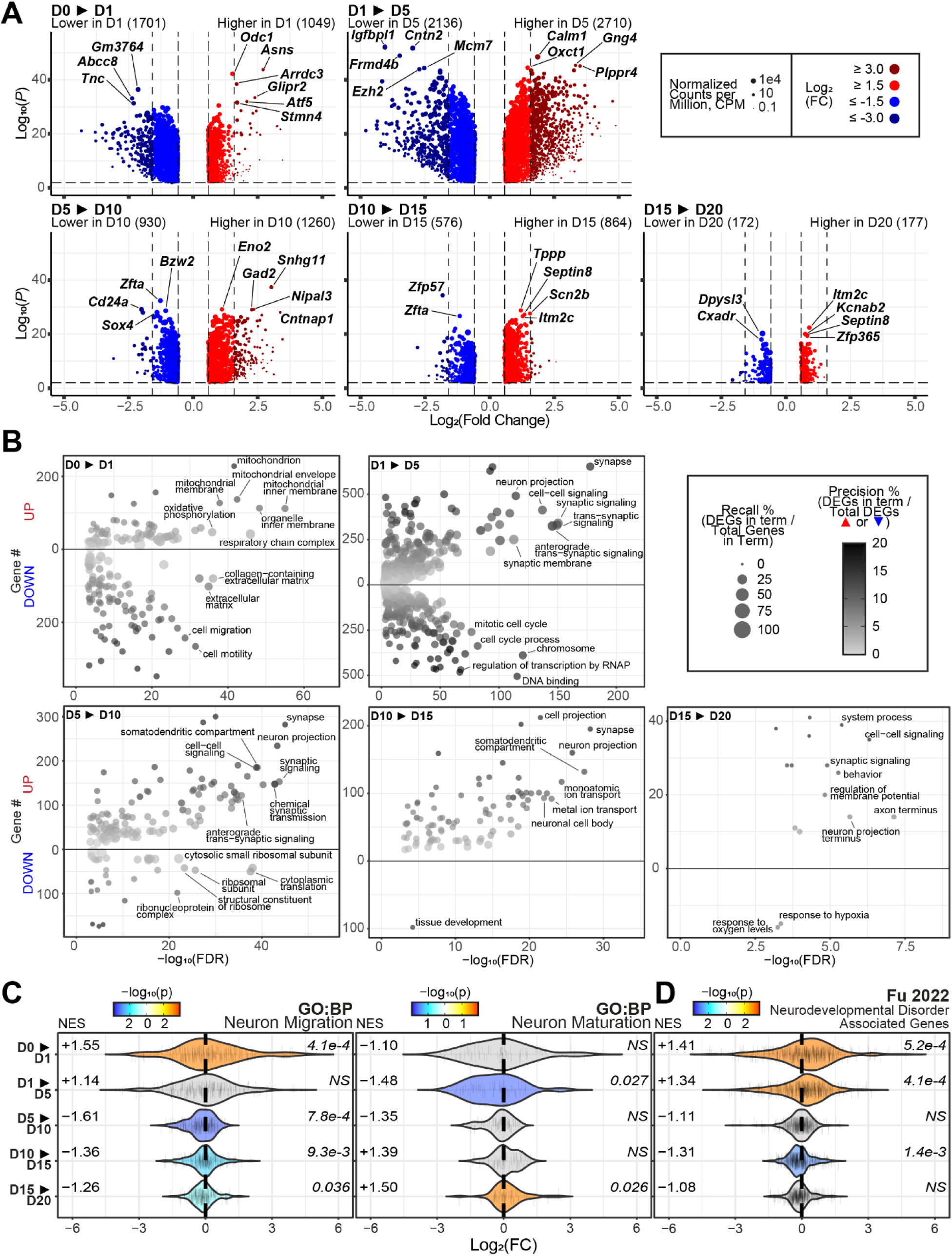
Stepwise differential gene expression analysis between consecutive time points across neuronal maturation. A. Volcano plots of DEGs from the pairwise comparisons of each time point and its subsequent time point. Points are scaled in size proportional to its CPM and colored by fold change. B. Gene ontology analysis of significantly upregulated and downregulated genes for each pairwise comparison. Only terms with up to 2000 genes were included. Points are scaled in size proportional to recall (the number of DEGs found per term against the total number of genes in that term) and colored by precision (the number of DEGs found per term against the total number of upregulated or downregulated DEGs used as the query). C. Violin plots of gene set enrichment analysis of genes in the GO term for neuron migration, neuron maturation, or neurodevelopmental disorder-associated genes from Fu et al (FDR < 0.05). NES indicates the directional normalized enrichment score across all genes. Multiple testing corrected significance values are shown to the right of each plot.

Highly significant gene ontology enrichment was observed for each stepwise comparison (Fig. 3B). Corroborating the timepoint-specific analysis, the D0-D1 transition showed robust upregulation of genes encoding mitochondrial components. There was a corresponding reduction in transcripts associated with locomotion, morphogenesis, and cell motility. Neuron-associated GO term enrichment emerged from D1 onward. SynGO analysis, specific to proteins with known functions at the synapse, revealed consistent upregulation of genes involved in both the pre and post synapse, synaptic signaling, and organization between D1 to D15 (Supplementary Fig. 4B), consistent with an active period of maturation and synaptogenesis. As anticipated, genes associated with mitosis, translation, and development declined over neuronal maturation.

To elucidate regulatory mechanisms underlying the transcriptional profiles, we performed transcription factor motif enrichment of the upstream promoter region of each stepwise DEG group (Supplemental Fig. 4C). We found multiple significantly enriched motifs, primarily among promoters of downregulated genes, including *Ngfr* and *Notch3*.

Because gene ontology analysis is dependent on fold change and significance thresholds that may overlook subtle but coordinated changes, we also performed gene set enrichment analysis of select transcript groups (Fig. 3C, Supplemental Fig. 4D). We examined genes within GO biological processes related to neuron migration and detected enrichment between 0 and 1 DIV and subsequent decreased expression after 5 DIV. Conversely, neuronal maturation genes show an opposing pattern with decreased expression between 1 and 5 DIV and increasing expression after 15 DIV. Using a single-cell atlas characterizing gene expression of developing excitatory neurons^51^, we also identified similar late enrichment patterns after D10 to D15. Additionally, we found that genes associated with neurodevelopmental^52^ increase between D0 to D5 than stabilize and modestly decrease after 10 DIV (Fig. 3D). Similar but non-significant trends are observed in genes specifically associated with autism (Supplemental Fig. 4D). Together, these findings delineate distinct phases of maturation captured throughout time *in vitro* for primary cultured neurons, reflecting both synapse development and neuronal maturation.

### Trajectory analysis of transcripts across development

To characterize genes exhibiting temporal changes across the full longitudinal period, we applied a regression model incorporating time and sex and clustering transcripts by shared trajectories. Unsupervised clustering confidently identified 7 linear (Fig. 4A) and 6 quadratic (Supplemental Fig. 4) trajectories. Linear clusters 1, 2, and 3 included transcripts consistently downregulated across days *in vitro* while clusters 5, 6, and 7 showed consistent upregulation (Fig. 4B).

**Figure 4:**
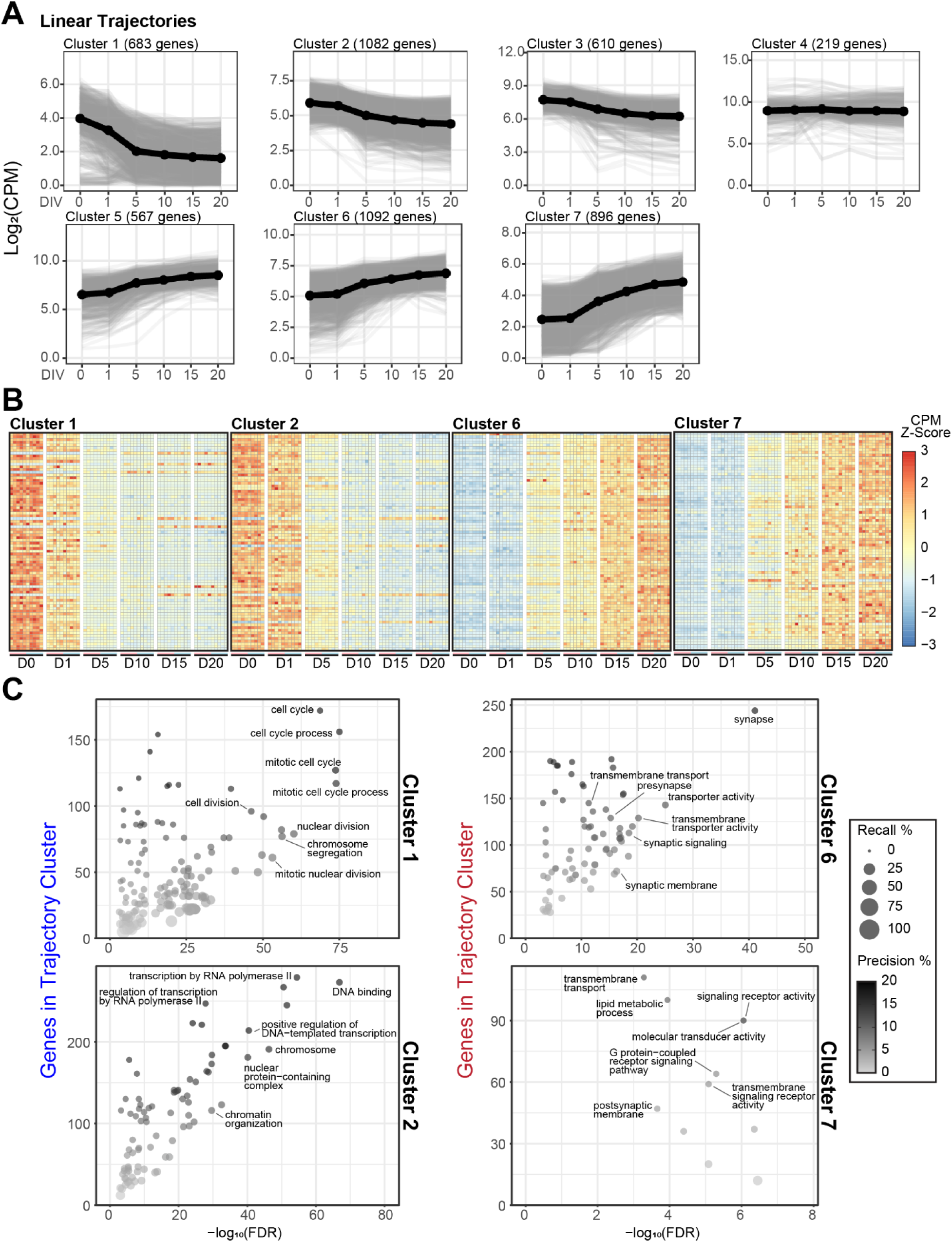
Longitudinal regression analysis of transcriptomic trajectories. A. Line plots of the CPM of all transcripts with significant linear trajectories as computed by maSigPro. The solid black line is the average CPM of all genes within a cluster. B. Heatmap of gene expression across time points for 100 randomly selected genes from 4 linear trajectory clusters. C. Gene ontology analysis of genes from 4 linear trajectory clusters.

Longitudinal trends recapitulated findings from the pairwise comparisons, including the consistent upregulation of synapse-associated transcripts (Fig. 4C) but also revealed changes beyond those observed in pairwise analysis. Two clusters demonstrated sustained downregulation from D0 to D20, comprising genes involved in cell cycle and transcriptional regulation. Among these were critical epigenetic regulators of neuronal differentiation such as *Sox2*^53,54^, *Pax6*^55,56^, and *Prox1*^57^. These findings support a model in which the *in vitro* system transitions from a more proliferative state to a specialized, non-dividing neuronal identity with a unique transcriptomic landscape.

Several clusters, such as linear clusters 1 and 7 and quadratic clusters 2, 3 and 5, exhibited more pronounced early slopes between DIV0-DIV1 than between time points. These differences likely reflect shifts in cell composition and adaptation to culture conditions during the earliest stages. However, many clusters demonstrated stable directional changes following these early time points, indicating a substantial number of genes that continue to either increase or decrease as neurons mature (Fig. 4A). Notably, few genes remained stable across this trajectory, with only linear cluster 4 (219 genes) showed modest variation whereas most clusters had highly dynamic gene expression changes in both linear and quadratic models (Supplemental Fig 5).

Gene ontology analysis of consistently downregulated clusters again showed a gradual decline in genes associated with mitosis, cell cycle, and transcription. Conversely, synapse-related genes and transmembrane transporters accounted for a significant proportion of upregulated clusters (Fig. 4C). No clusters displayed a mid-trajectory peak in expression, likely reflecting the continuous maturation of neurons during this period and the stable postmitotic identity established shortly after DIV1 (Supplemental Figure 1).

### Sex-specific differential gene expression during neuronal maturation

This dataset included 5 female and 5 male replicates for each time point, enabling us to assess sex-specific transcripts. Other than the known sex-specific genes such as X-linked genes that escape X-inactivation (*Xist*, *Kdm5c*, *Kdm6a*, *Ddx3x*, *Eif2s3x*) or those on the Y-chromosome (*Kdm5d*, *Uty*, *Uba1y, Ddx3y*, *Eif2s3y*), we found no significant DEGs in any timepoint-specific or stepwise comparison. Differences were found, however, in the longitudinal regression model factor for sex (Supplemental Fig. 6A). Unbiased clustering identified clusters with gene sets exhibiting slight deviations in expression between female and male mouse cortices that persisted for only one or two days. There were no significant gene ontology or protein family patterns among these clustered genes.

Regression also identified 7 transcripts that represented their own independent clusters (Fig. 5A). Among those, *Cort*, *Tac1*, and *Oip5os1* showed a greater increase in expression from D10 through D20 in females than males. Both *Tac1* and *Cort,* encode neuropeptides, Neurokinin A and Cortistatin, respectively. Differences in *Cort1* were particularly robust and detectable even without trajectory analysis (Fig. 5B) and validated using qPCR in an independent set of cultured neurons (Supplemental Fig. 6B). Interestingly, all of these sex-specific differences emerged at late time points, well after cells have been isolated from *in vivo* signals and hormonal inputs. This suggests that they may reflect cell-intrinsic differences in genome regulation between male and female mouse neurons, potentially downstream of sex-chromosome gene effects rather than a response to extracellular cues.

**Figure 5:**
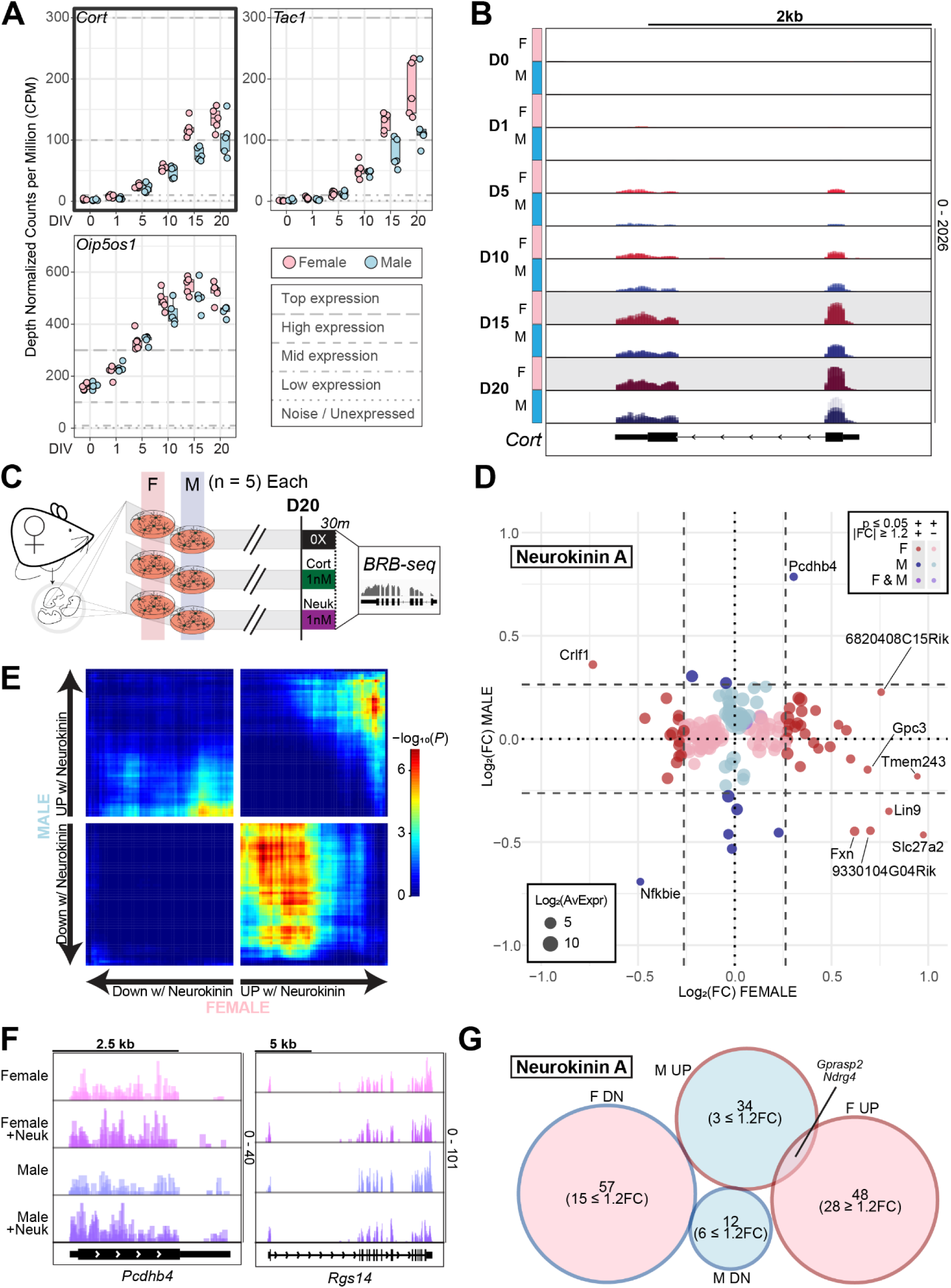
Sex-specific neuronal maturation transcripts. A. Counts per million (CPM) box plots of the expression of sex-specific transcripts over days *in vitro* (n = 5 per sex). B. Gene track of aligned transcripts over the female-specific temporally upregulated gene *Cort*. Replicates by time and sex are overlayed for each track. C. Schematic of the experimental timeline for the comparison of transcriptomes of female- and male-derived neurons at DIV20 treated with either water vehicle, 1nM Cortistatin, or 1nM Neurokinin A for 30 minutes. D. Scatterplot of the fold changes of genes differentially expressed after Neurokinin A treatment in either male or female neurons. Only those genes with adjusted(*p*) ≤ 0.05 in at least one sex are shown. E. Rank–Rank Hypergeometric Overlap of adjusted(*p*) ranked genes between female or male Neurokinin A treatment. F. Gene tracks of aligned transcripts over select, sex-specific DEGs in the Neurokinin A treatment. G. Venn diagram of the overlap of neurokinin A treatment DEGs between sexes and directionality.

To determine whether sex-specific differences in precursor expression translated into broader differences in neuropeptide signaling, DIV20 male and female-derived neurons were treated with either cortistatin-14 or neurokinin A for 30 minutes before RNA collection and multiplexed, bulk RNA-sequencing (Fig. 5C). Cortistatin caused very few gene expression changes, potentially because higher basal levels may occlude treatment effects (Supplemental Fig. 6C-D). In contrast, neurokinin A treatment resulted in modest but almost entirely sex-specific expression changes (Fig. 5D). Female neurons exhibited both the greatest number of significant DEGs, and greater fold changes. To determine if neuropeptide-induced expression changes were distinct across all genes between the sexes beyond DEGs identified by edgeR, we performed Rank-Rank Hypergeometric Overlap (RRHO) for each treatment (Fig. 5E, Supplemental Fig. 6E). We found that the Cortistatin response was convergent between sexes, while Neurokinin A treatment drove dramatically divergent male and female responses. There was no detectable concordance amongst Neurokinin A downregulated transcripts. Amongst upregulated transcripts, the most Neurokinin A-responsive genes were similar, but those with moderate upregulation in female neurons were downregulated in males. Although no functional groups were enriched amongst the male or female responsive genes, we found numerous cell adhesion components such as *Pcdhb4* (up in males), *Pdlim1* (down in females), and *Flna* (up in females) and unique responses in important signaling molecules such as *Rsg14* (down in males) (Fig. 5F). When examining DEGs that met significance criteria, we found that female and male neurons shared almost no Neurokinin-A responsive transcripts except for upregulation of the G protein-coupled receptor regulator *Gprasp2* and alpha/beta hydrolase *Ndrg4* (Fig. 5G).

Together, these results represent, to the best of our knowledge, the first identification of primary neuron culture sex-specific differences in autosomal gene expression that emerge upon neuronal maturation even in the absence of *in vivo* hormonal signals. They also reveal previously unrecognized neuropeptide-specific responses in cortical neurons, including striking sex-dependent patterns.

## Discussion

Here, we compare the gene expression profiles of primary mouse cortical neuron cultures at six time points *in vitro* using RNA sequencing. Primary neuron cultures exhibit expected changes in the expression of key genes involved in neuronal development *in vivo*, supporting the biological relevance of the model. Stage-specific, consecutive pairwise, and longitudinal regression analyses revealed transcripts and transcriptomic signatures that change across neuronal maturation. Further, by independently assessing neurons derived from female and male brains we unexpectedly identified sex-specific expression trajectories for key neuropeptides. The dataset generated here provides a valuable resource for researchers utilizing mouse cortical neuron cultures or comparing the culture to *in vivo* neuronal maturation models. Notably, we also identified the first sex differences in gene expression, beyond those linked to sex chromosomes, that are detectable in neuronal culture systems even in the absence of external influences.

Within the many gene expression changes described here, continued upregulation of neuron, synapse, and synaptic signaling components is consistent across all subsequent days in culture (Fig. 3B, Supplemental Fig. 3B). Day-specific analysis comparing each time point against the aggregate of the others recapitulated the general observation from principal component analysis that the earliest time points, particularly D0 which precedes culturing, harbor the most unique DEGs. However, also noteworthy was the emergence of time-point-specific DEGs at DIV20, particularly those involved in neuronal signaling responses. We and others have shown that mouse cortical cultures exhibit spontaneous network firing activity from D18 onward^58,59,19^. These transcriptomic changes may provide insights into the gene expression programs linked to functional maturation apparent and network activity.

A particularly exciting finding to emerge from this dataset is that neuropeptide-encoding such as *Cort* and potentially *Tac1* increase in expression over maturation time to a greater degree in neurons derived from female tissue. Tachykinins, the active products generated by the *Tac1* precursor, include Substance P and Neurokinin A, and are critical in and beyond the nervous system^60,61^. Substance P and Neurokinin A are essential in the inflammatory pain response^62,63^. Although differences in pain response by sex are well documented^64,65^, transcriptional regulatory processes that control their expression in cortical neurons, particularly during development and maturation, as well as the functions of *Tac1* products, are not well understood. *Cort* encodes Cortistatin, which is structurally similar to somatostatin (*Sst*). Both are involved in synaptic plasticity^66^ and are essential in sexual dimorphism in fasting-induced pituitary growth hormone responses^67,68^. These neuropeptides play roles in the excitatory-inhibitory balance in the cortex but, similar to *Tac1*, their functions across maturation, and how they may differ between the sexes, remain unexplored.

The ability to detect sex differences in neuropeptide expression in primary neuron cultures, several weeks removed from the presence of hormones and other environmental effects, is remarkable. Because these neuropeptide genes are not on sex chromosomes, their differential expression suggests the presence of regulatory elements influenced by X or Y-linked factors. For example, *Kdm5c* is an X-linked gene that escapes X-inactivation and functions as a histone demethylase modulating gene expression. However, whether *Kdm5c* or other X or Y-linked genes can directly regulate neuropeptide expression remains unknown. Notably, these expression differences only clearly emerged *after* multiple weeks in culture. This suggests that cortical neurons contain innate regulatory mechanisms that lead to sex-dependent expression patterns of neuropeptides. Prior work has extensively examined sexual dimorphism in behavior^69–71^ and development^72^. These studies include RNA-sequencing in mouse^73^ and human^74,75^ embryonic stem cells through differentiation into mature tissues^76,77^. Investigations into sex-differences consistently identify subtle but reproducible differences in autosomal gene expression related to cell fate determination and epigenetic regulation. Our findings add to these findings by capturing a previously unresolved aspect of sex-specific biology occurring between neuron differentiation and maturity. We also provide this data in a easily viewable and downloadable format to allow researchers to quickly check genes of interest to identify potential sex differences in expression and relevant time points at which to perform experiments.

Despite the strength of our dataset, the time course has several inherent limitations. Variability between individual embryos introduces substantial biological noise. While 10 biological replicates (or 5 per sex per time) provided sufficient statistical strength to detect numerous changes, rare genes or differences with small effect sizes may remain undetected. Furthermore, our sequencing strategy of poly-A enrichment with short-read libraries of 10–15 million reads per sample imposes constraints. Such parameters limit detection of non-polyadenylated transcripts, low-abundance RNAs, and lack power for intensive isoform analysis. Emerging long-read sequencing with greater depth and isoform resolution would overcome these limitations and may further clarify the sex-specific transcriptional patterns. Finally, although primary neuron cultures provide a powerful and tractable model for mechanistic studies, they cannot fully recapitulate the structural complexity, circuit architecture, and multicellular interactions present *in vivo*.

In summary, we generated a comprehensive longitudinal RNA-seq dataset of primary cortical neuronal cultures derived from embryonic mice. These results revealed extensive transcriptomic changes across maturation and uncovered previously unrecognized sex-specific expression patterns in maturing neurons. These datasets offer valuable resources and references for researchers using primary neuronal cultures and highlights important distinctions between *in vitro* and *in vivo* systems. Moreover, these findings may provide key insights into the regulation and functional significance of sex-specific neuropeptide pathways in the cortex.

## Methods

### Primary cell culture

Cortices were dissected from wildtype E16.5 C57BL/6J embryos and cultured in neurobasal medium (Gibco 21103049) supplemented with B27 (Gibco 17504044), GlutaMAX (Gibco 35050061), penicillin-streptomycin (Gibco 15140122) in 12-well plates coated with 0.05 mg/mL Poly-D-lysine (Sigma-Aldrich A-003-E). Neurons were maintained at 5% CO_2_ and 37°c. Two litters at E16.5 were dissected in parallel, and all cortices were seeded. Both cortices of each individual pup were resuspended in 6mL of media, and 5mL were used to seed 5 wells per pup. At 3 DIV, neurons were treated with 0.5 µM AraC with additional media added (1X volume). The remaining 1mL was kept for the 0 DIV timepoint RNA collection. A half media change was performed at 10, 14, and 18 DIV. For the neuropeptide treatment experiments, 1mM solutions of each neuropeptide in water were diluted 1:1000 in the cell media. All experiments were performed in accordance and with approval of the University of Pennsylvania IACUC protocol number 806617.

### Genotyping for pup sex

Tissue was collected from the carcass of each pup during cortex dissection and denatured in 25mM NaOH, 0.2mM EDTA in water at 95°c for 1 hour. Denaturing was neutralized in 40mM Tris HCl (pH 5) in water. Genomic DNA was purified using the DNA Clean and Concentrator kit (Zymo Research). Sex was determined by PCR amplification of primers specific to the X and Y chromosome using the KAPA2G Fast HotStart PCR Kit (Roche). Primers are described in McFarlane et al^78^.

### Library preparation and sequencing

RNA was isolated using the Zymo Quick-RNA Miniprep Plus Kit (R1057). 200µL of lysis buffer was added to either the 1mL of remaining neurons following plate seeding (0 DIV) or from 1 well per pup at 1, 5, 10, 15, and 20 DIV. 5 female and 5 male samples per timepoint were selected based on the concentrations and quality of RNA across timepoints. 150ng of RNA per sample, per timepoint were processed using the standard protocol of the NEBNext Poly(A) mRNA Magnetic Isolation Module and then immediately loaded into 60 wells of the Mercurius BRB-seq library preparation kit (V5, Alithea Genomics 10801). All samples were pooled and processed in one sublibrary (MF.UDI.1) to minimize batch effects. Prior to sequencing, library size distribution was confirmed by capillary electrophoresis using an Agilent 4200 TapeStation with high sensitivity D1000 reagents (5067-5585), and libraries were quantified by qPCR using a KAPA Library Quantification Kit (Roche 07960140001). Libraries were sequenced on an Illumina Nextseq2000 P3 with 1% PhiX spike-in and a loading concentration of 850pM (20 million reads per sample, 90-bp read length).

### Raw data processing and demultiplexing

BCL files representing all 60 samples were converted into multiplexed fastq format using bcl2fastq (V2.20) in the Illumina BaseSpace Sequence Hub. FastQC (V0.11.2) was performed on the R2 file corresponding to the cDNA reads. STAR solo (V2.7.8a) was used to create the raw count matrices using the following parameters: --ruMode alignReads, --outSAMmapqUnique 60, --quantMode GeneCounts, --outFilterMatchNminOverLread 0.51, --soloType CB_UMI_Simple, --soloUMIstart 15, --soloUMIdedup NoDedup 1MM_Directional, --soloCellFilter None, --soloFeatures Gene, --soloAdapterMismatchesNmax 2, --outFilterMultimapNmax 20. Samples were demultiplexed in STAR using the barcodes_v5C_96_star.txt file provided by Alithea Genomics specifying the index information. Reads were aligned and assigned to genomic features using the mm10 genome. Individual bam files for each sample were generated and indexed using samtools (V1.1). Deeptools (V3.5.6) bamCoverage was used to generate bigwig files with a sliding window of 20 and RPKM normalization.

### Data normalization

For analyses involving all timepoints, or one timepoint against all other timepoints, EdgeR (V4.4.1) was used for normalization and model fitting. Normalization factors were calculated with the trimmed mean of M-values method using the upper quantile of counts and the asymptotic binomial precision weights. Samples were scaled by the normalization factors to convert observed library sizes to normalized library sizes for all subsequent steps. Genes with a minimum normalized count over a normalized counts per million (CPM) of 1 were filtered prior to transformation.

### Differential gene expression analysis

Count data was transformed using voom and duplicate correlation was performed with the subsequent vGene matrix, using the sex:time IDs as the blocking variable. Model fitting was performed using lmFit specifying the consensus correlation between samples and the same blocking variable. Finally, t and F statistics were calculated using empirical bayes statistics. Stepwise comparisons were made using makeContrasts for (Time(n) – Time (n-1)), where n corresponds to each sequential time point. “1vAll” comparisons were made using makeContrasts for one time point subtracted by the average of all other 5 timepoints. All differential expression analysis between time points used 5 female and 5 male samples as replicates. Differentially expressed genes (DEGs) were specified as follows: baseMean of CPM > 1, adjusted p value ≤ 0.05, and |fold change| ≥ 1.5 (|log2(FC)| ≥ 0.585).

### Downstream analysis

Imputation of bulk admixture expression identities from single-nuclei data was performed using CIBERSORTx^79^. CPM normalized counts were used for each time point. The single-nuclei signature matrix was generated using the cluster identity labels for P40 mouse prefrontal cortex isolated nuclei (GSE298627). Average expression values were calculated for each identity-labelled nuclei.

Gene ontology analysis was performed using gProfiler2^80^ (v0.2.3) over-representation analysis against the GO molecular function, cellular component, and biological process databases. Multiple testing was corrected using the g:SCS algorithm, a GO-optimized method that provides a better threshold between significant and non-significant than FDR^81^. Only terms with ≥ genes were chosen for specificity. The domain scope of GO testing used a background list of all genes expressed in at least 1 time point (for the 1vAll comparisons), or all genes expressed in either time point n or n-1 (for the stepwise comparisons). In both cases, only genes with a baseMean of normalized counts > 1 were used. SynGO was performed using the web client^82^ (v1.2), specifying a medium stringency which only includes annotations that also use biological systems that are not cell-free, and do not solely rely on biochemical fractionation.

Transcription factor motif analysis on 5kb windows directly upstream of stepwise DEGs were analyzed using sequence enrichment analysis in the MEME suite^83^ (v5.5.7). Motifs were scanned using the JASPAR 2020 Core Vertebrates database using the windows 5kb upstream of 1000 randomly selected genes expressed in neurons.

Gene set enrichment analysis was performed using fGSEA^84^ (V1.32.2) with lists from the gene ontology database, the Descartes organogenesis single-cell transcriptional atlas^51^, and the cell-type-specific gene expression atlas of the mouse neocortex^85^.

Temporal expression changes and differences between female and male neurons were assessed using maSigPro^86^. Each 5 mice per time point were assigned as replicates to two either a female or male condition in the design matrix. The matrix of CPM normalized counts across all time points was assigned to a quadratic regression and fit using the p.vector function with the following parameters: Q = 0.05, MT.adjust = “BH”, min.obs = 1, theta = 10. The best fit regression model for each gene was selected using T.fit with multiple testing correction and a two-way backwards step procedure for all variables. Significant genes exhibiting linear, quadratic, or male-biased trajectories were determined using see.genes with the optimal number of clusters competed using the Mclust algorithm and multiple testing correction.

Rank–Rank Hypergeometric Overlap (RRHO) was performed using the RRHO2 package^87^. DEG lists were rank-ordered by multiplying the adjusted *p* against the direction of the fold change and then merged. Significance was calculated using the hypergeometric test with BH correction.

### RT-qPCR

500ng of RNA per sample was used to prepare cDNA using the high-capacity cDNA reverse transcription kit (Applied Biosystems 4368813), and quantitative PCR was performed with Power SYBR Green PCR master mix (Applied Biosystems 4367659).

*Cort*: F- GGTCGCAGCCTCCGCCCTTC, R- TTGGGAAGCCCACTCGTGCCA.
*Gapdh*: F- AACTCACTCAAGATTGTCAGCAA, R- GGCATGGACTGTGGTCATGA

### Statistical analysis

All statistical analysis were performed using readily available code in R. The number of replicates and details of statistical tests are reported in figure legends and methods.

## Author Contributions

Conceptualization: AP, EK; methodology: AP, EK; investigation: AP, RA; visualization: AP; formal Analysis: AP; website development: JG; supervision: EK; writing – original draft: AP; writing – review and editing: AP, EK; funding acquisition: EK.

## Data and code availability

The processed time course data are available for download and visualization at https://www.korblab.com/data. RNA-sequencing and single nucleus RNA-sequencing data generated in this study will be available under the following GEO accession number GSE324205 upon publication. All data are available in the main text or the supplementary materials. Any information required to reanalyze the data reported in the paper is available from the lead contact upon request.

## Lead contact

Requests for further information, resources, and reagents should be directed to and will be fulfilled by the lead contact, Erica Korb.

## Materials availability

This study did not generate new, unique reagents.

---

**Supplemental Figure 1:**
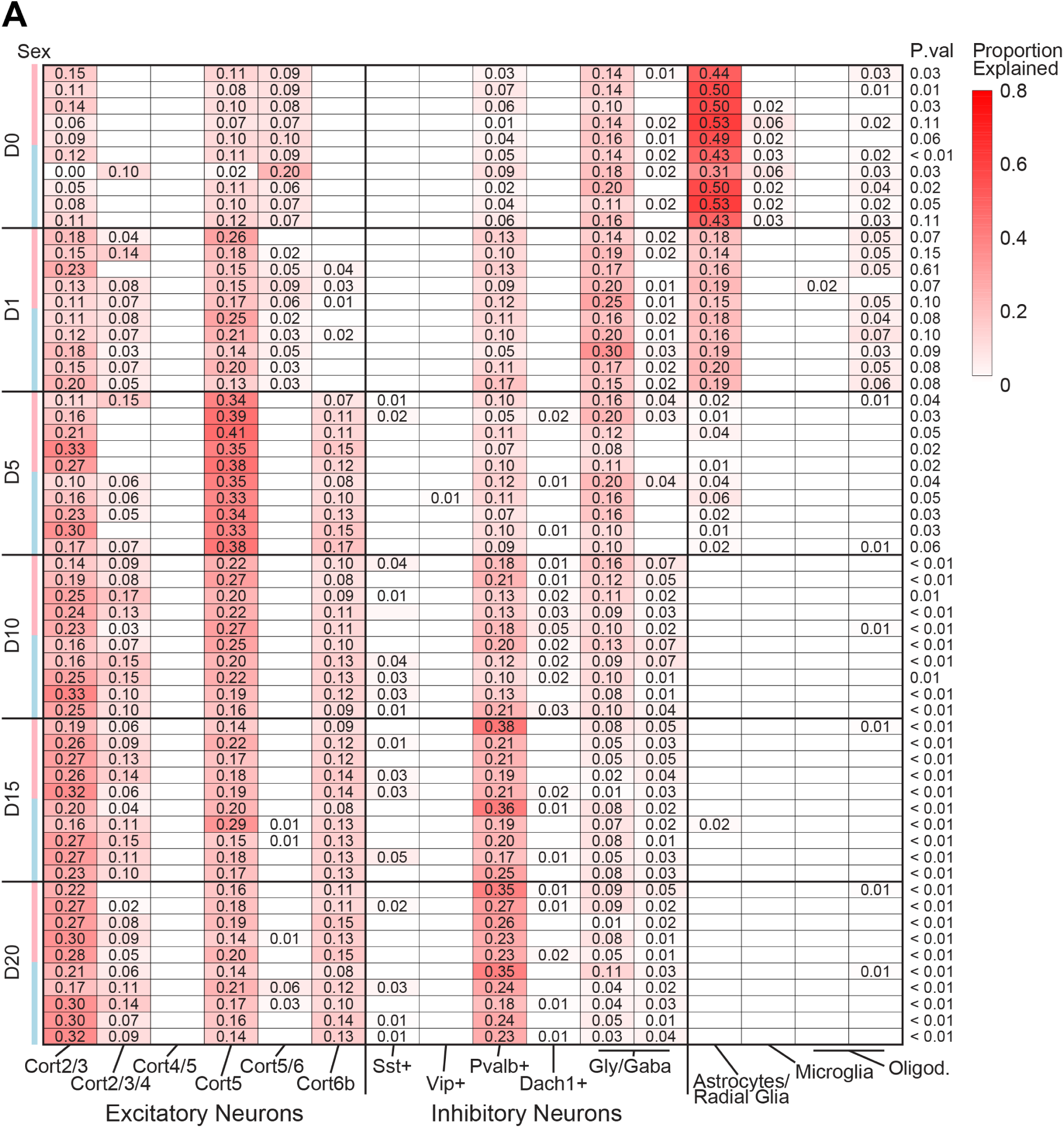
Deconvolution of bulk transcriptomes. A. Proportion testing of each replicate bulk gene expression profiles using cell population cluster assignments from a wildtype P40 mouse cortex single-nuclei dataset. Permutation testing of the bulk profile imputation was performed for each sample for the significance (p) value.

**Supplemental Figure 2:**
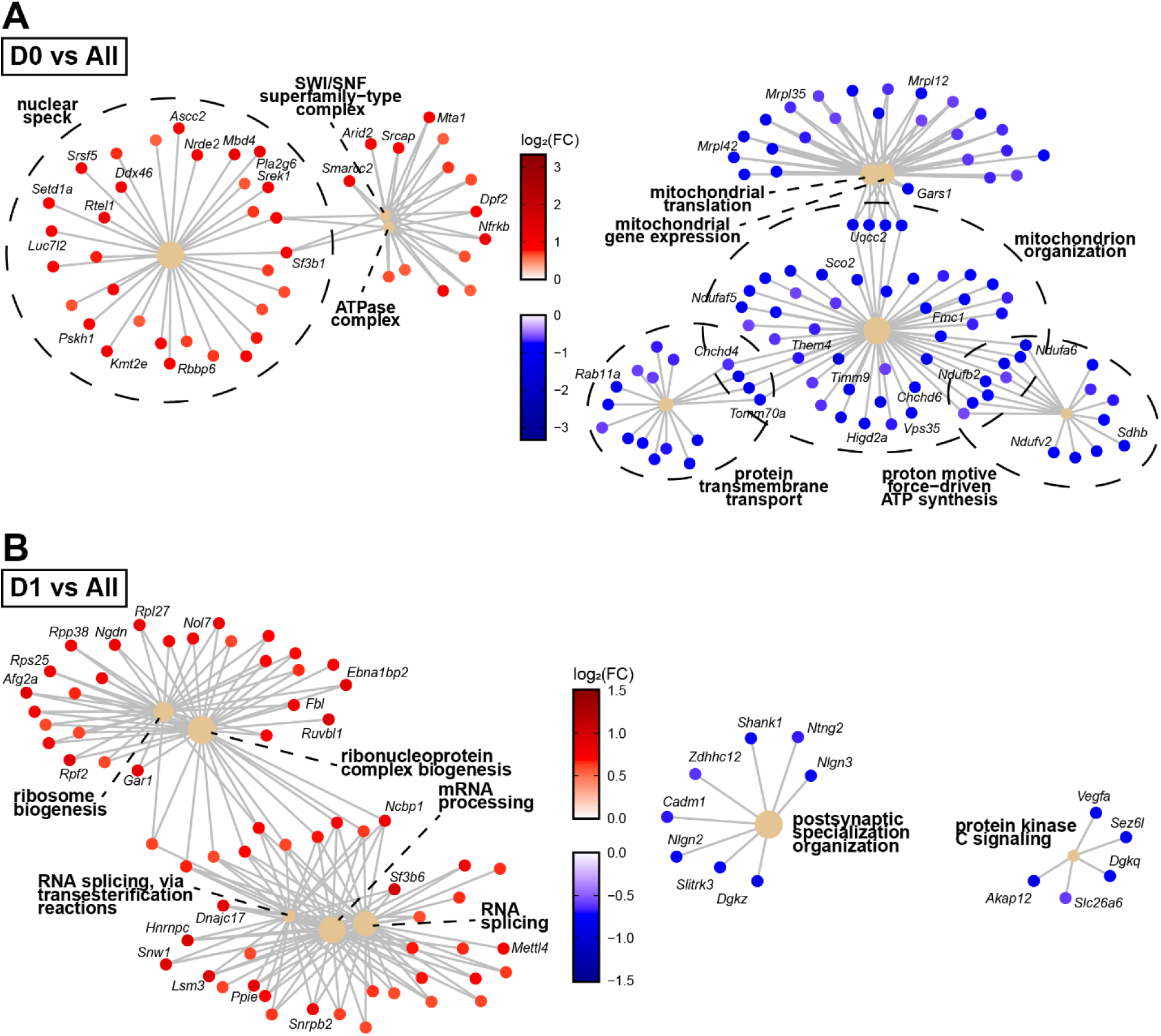
Gene ontology analysis of timepoint-specific DEGs. A-B. Category-Network plots of gene ontology analysis of significantly upregulated and downregulated genes for the comparison between D0 (A) or D1 (B) against the regression aggregate of all other time points. Only those terms with a multiple testing corrected FDR ≤ 0.05 were shown for clustering.

**Supplemental Figure 3:**
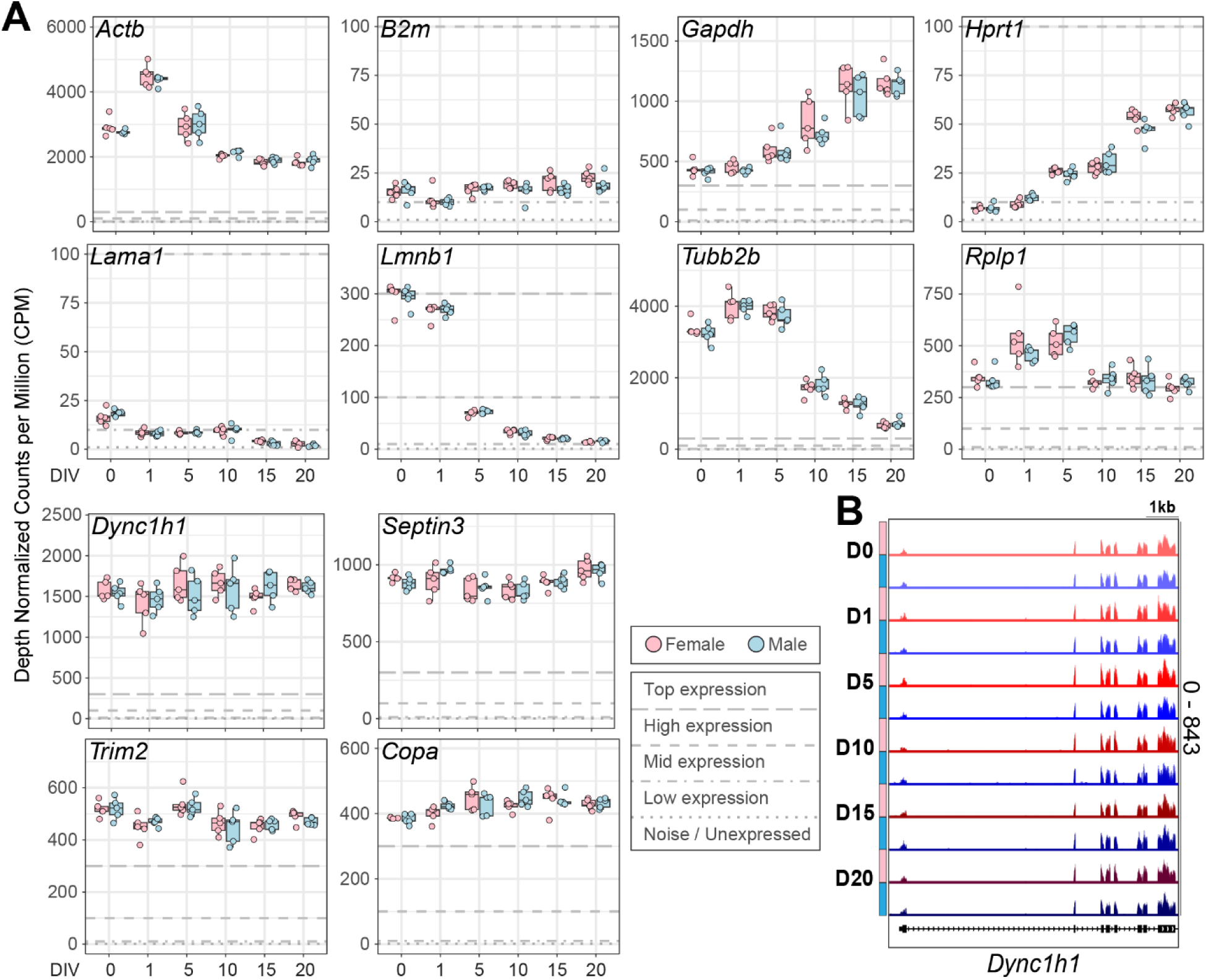
Temporal expression of housekeeping genes. A. Counts per million (CPM) box plots of the expression of known and novel housekeeping genes over days *in vitro* (n = 5 per sex). Bolded lines demarcate thresholds of very high to high (CPM > 300), high to mid (300 > CPM > 100), mid to low (100 > CPM > 10), and low to noise limit expression (10 > CPM). B. Gene track of aligned transcripts over the 5’ end of the *Dync1h1* gene.

**Supplemental Figure 4:**
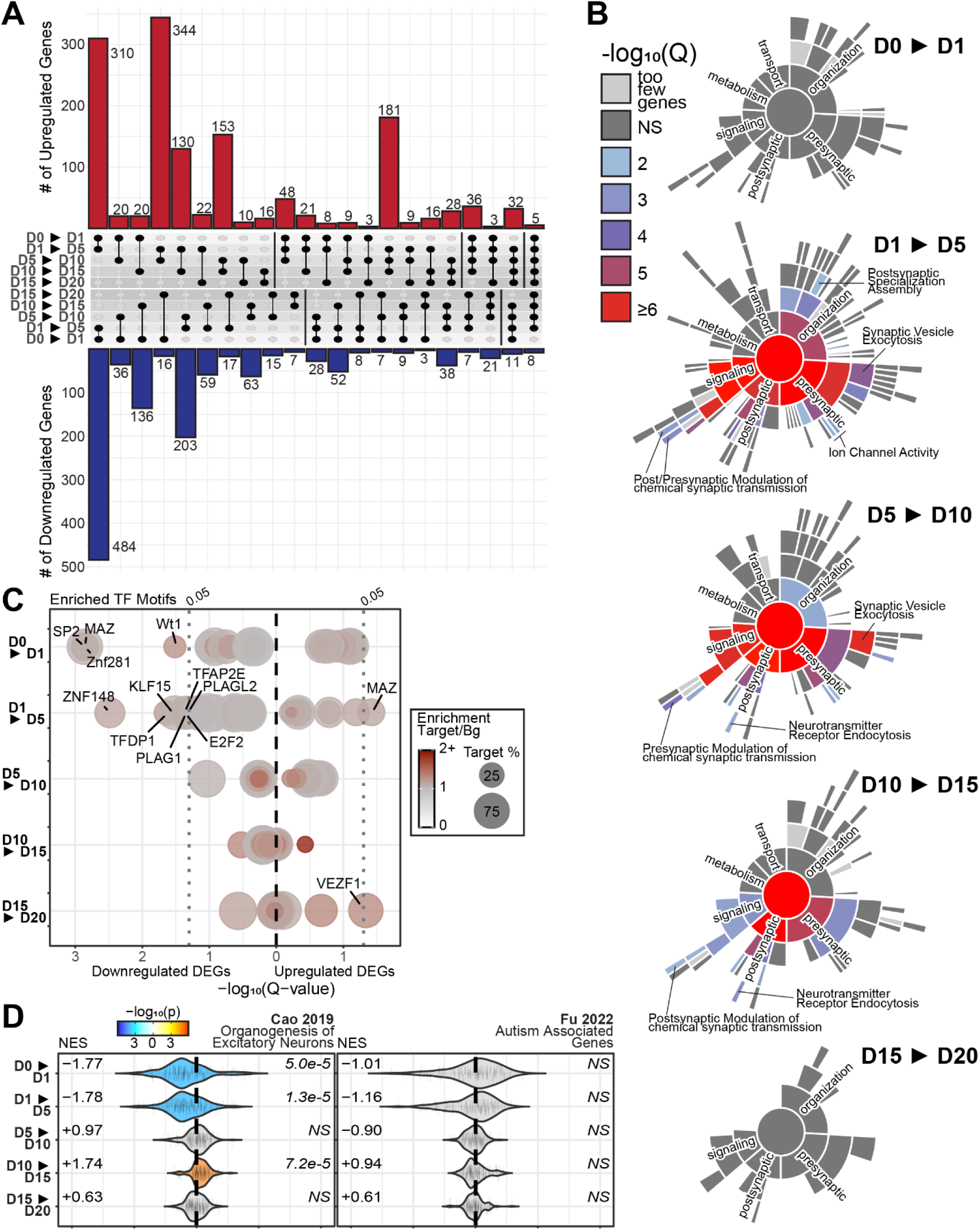
Intersection and ontology analysis of stepwise DEGs. A. Upset plot of up or downregulated DEGs found in at least pairwise time point comparisons. Gene placement prioritizes the largest intersection number possible. B. Synapse-specific gene ontology (SynGO) analysis of significantly upregulated genes in each pairwise time point comparison. Ontology hyponyms with too few genes were omitted from visualization unless they were an umbrella term for at least one hyponym with enough genes for significance testing. C. Transcription factor motif enrichment analysis of the 5kb regions upstream of all significant DEGs by time point transition. Points are scaled in size proportional to the number of DEG promoter regions containing the motif and colored by the enrichment proportion of that motif in DEGs versus all genes expressed in either time point in the comparison. D. Violin plots of gene set enrichment analysis of marker genes for developing excitatory neurons or autism associated genes from Fu et al (FDR < 0.05). NES indicates the directional normalized enrichment score across all genes. Multiple testing corrected significance values are shown to the right of each plot.

**Supplemental Figure 5:**
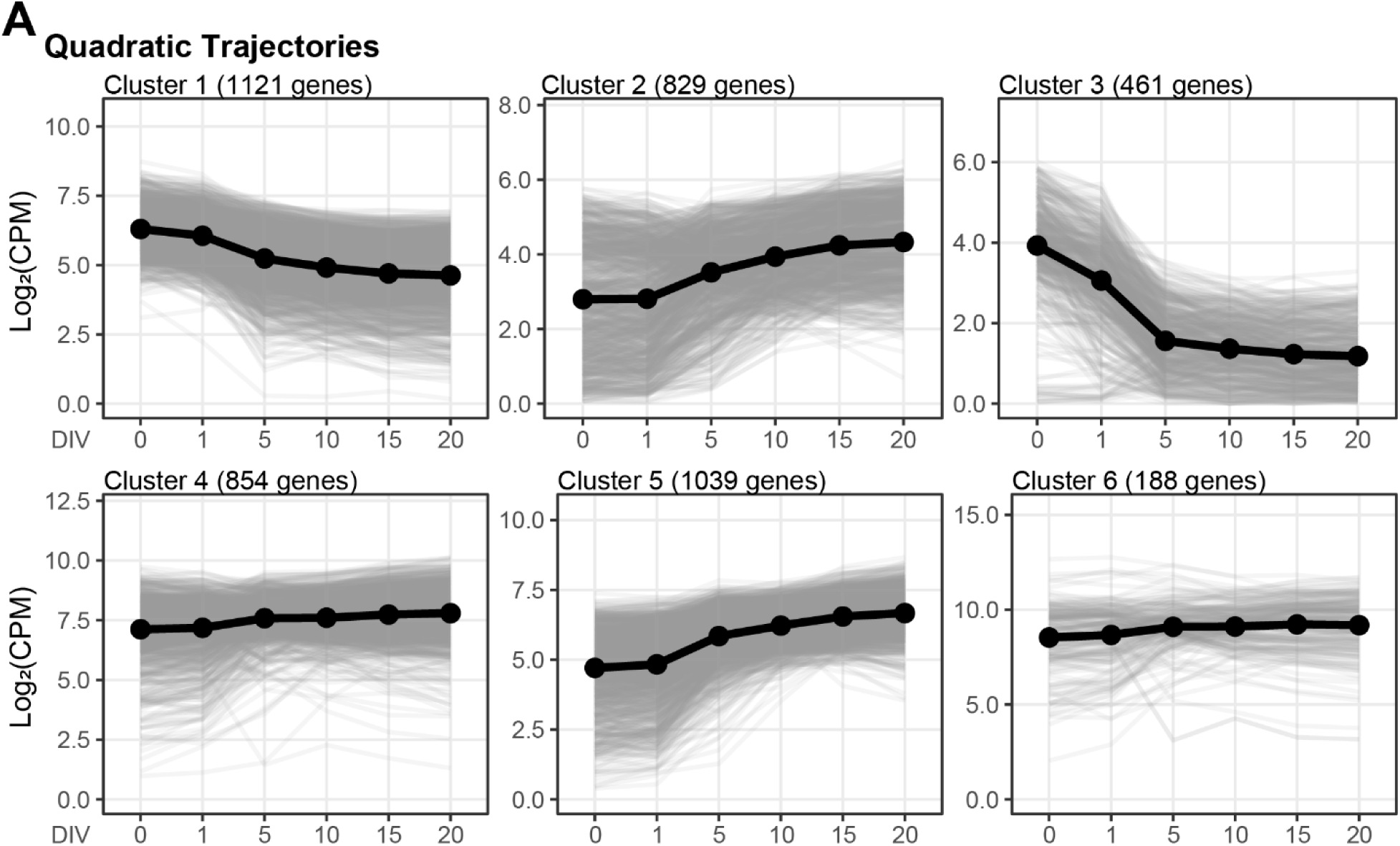
Quadratic trajectories. A. Line plots of the CPM of all transcripts with significant quadratic trajectories as computed by maSigPro. The solid black line is the average CPM of all genes within a cluster.

**Supplemental Figure 6:**
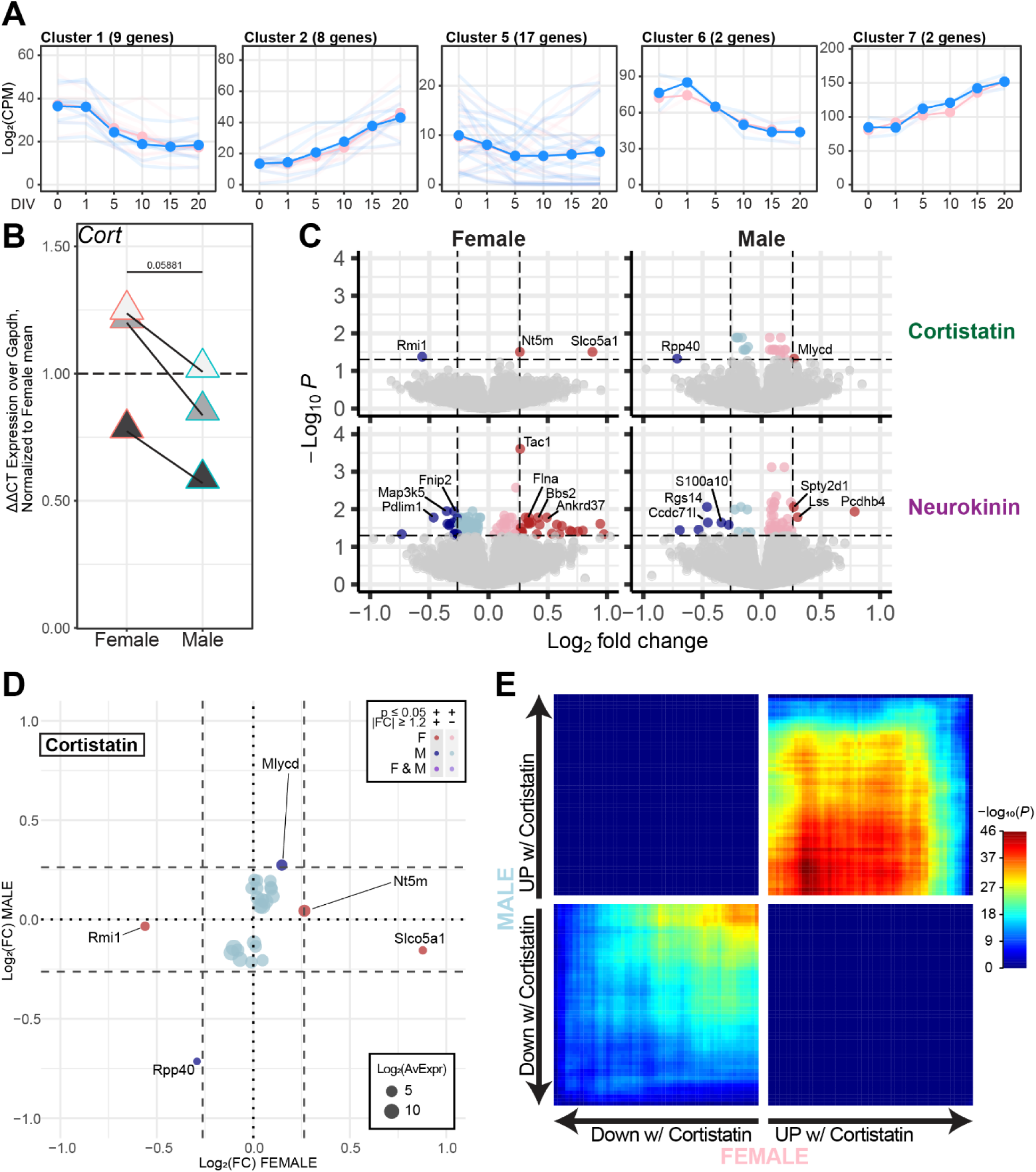
Sex-specific trajectory modules and neuropeptide treatment. A. Line plots of the CPM of all transcripts with significant male-deviated trajectories containing at least 2 genes per cluster, as computed by maSigPro. Faint lines representing the individual gene trajectories or solid lines for the average of genes within the clusters are colored pink (female) or blue (male). B. Relative transcript expression level of *Cort* as measured by RT-qPCR ΔΔCT over *Gapdh* (n = 5 per sex). Samples are plotted relative to the average of the female neurons and colored by litter of origin. C. Volcano plots of differentially expressed genes in either female or male neurons following 30 minute treatment with either Cortistatin or Neurokinin A. DEGs up or downregulated with a fold change ≥ 1.2 are in dark red or dark blue, respectively. D. Scatterplot of the fold changes of genes differentially expressed after Cortistatin treatment in either male or female neurons. Only those genes with adjusted(*p*) ≤ 0.05 in at least one sex are shown. E. Rank–Rank Hypergeometric Overlap of adjusted(*p*) ranked genes between female or male Cortistatin treatment.

## References

1. Banker, G. A. & Cowan, W. M. Rat hippocampal neurons in dispersed cell culture. Brain Res 126, 397–342 (1977).

2. Dichter, M. A. Rat cortical neurons in cell culture: culture methods, cell morphology, electrophysiology, and synapse formation. Brain Res 149, 279–293 (1978).

3. Weiss, S., Sebben, M. & Bockaert, J. Corticotropin-Peptide Regulation of Intracellular Cyclic AMP Production in Cortical Neurons in Primary Culture. Journal of Neurochemistry 45, 869–874 (1985).

4. Stevens, T. R., Krueger, S. R., Fitzsimonds, R. M. & Picciotto, M. R. Neuroprotection by Nicotine in Mouse Primary Cortical Cultures Involves Activation of Calcineurin and L-Type Calcium Channel Inactivation. J. Neurosci. 23, 10093–10099 (2003).

5. Monnerie, H. & Le Roux, P. D. Glutamate receptor agonist kainate enhances primary dendrite number and length from immature mouse cortical neurons in vitro. J of Neuroscience Research 83, 944–956 (2006).

6. Grabrucker, A., Vaida, B., Bockmann, J. & Boeckers, T. M. Synaptogenesis of hippocampal neurons in primary cell culture. Cell Tissue Res 338, 333–341 (2009).

7. Kim, S. H., Lu, H. F. & Alano, C. C. Neuronal Sirt3 Protects against Excitotoxic Injury in Mouse Cortical Neuron Culture. PLoS ONE 6, e14731 (2011).

8. Seibenhener, M. L. & Wooten, M. W. Isolation and Culture of Hippocampal Neurons from Prenatal Mice. JoVE 3634 (2012) doi:10.3791/3634.

9. Yang, L. et al. Neuroprotective effects of vitexin by inhibition of NMDA receptors in primary cultures of mouse cerebral cortical neurons. Mol Cell Biochem 386, 251–258 (2014).

10. Chen, W. et al. Tumor necrosis factor-α enhances voltage-gated Na+ currents in primary culture of mouse cortical neurons. J Neuroinflammation 12, 126 (2015).

11. Tyssowski, K. M. et al. Different Neuronal Activity Patterns Induce Different Gene Expression Programs. Neuron 98, 530–546.e11 (2018).

12. Dong, Y., Lozinski, B. M., Silva, C. & Yong, V. W. Studying the microglia response to oxidized phosphatidylcholine in primary mouse neuron culture and mouse spinal cord. STAR Protocols 2, 100853 (2021).

13. Chamberlin, S. R. et al. Multi-Omics Analysis in Mouse Primary Cortical Neurons Reveals Complex Positive and Negative Biological Interactions Between Constituent Compounds of Centella asiatica. Pharmaceuticals 18, 19 (2024).

14. Steiner, R. C., Heath, C. J. & Picciotto, M. R. Nicotine-induced phosphorylation of ERK in mouse primary cortical neurons: evidence for involvement of glutamatergic signaling and CaMKII. Journal of Neurochemistry 103, 666–678 (2007).

15. Fassier, C. et al. Microtubule-targeting drugs rescue axonal swellings in cortical neurons from spastin knock-out mice. Disease Models & Mechanisms dmm.008946 (2012) doi:10.1242/dmm.008946.

16. Ho, P. W.-L. et al. Age-dependent accumulation of oligomeric SNCA/α-synuclein from impaired degradation in mutant LRRK2 knockin mouse model of Parkinson disease: role for therapeutic activation of chaperone-mediated autophagy (CMA). Autophagy 16, 347–370 (2020).

17. Sun, Z., Williams, D. J., Xu, B. & Gogos, J. A. Altered function and maturation of primary cortical neurons from a 22q11.2 deletion mouse model of schizophrenia. Transl Psychiatry 8, 85 (2018).

18. Thudium, S., Palozola, K., L’Her, É. & Korb, E. Identification of a transcriptional signature found in multiple models of ASD and related disorders. Genome Res. 32, 1642–1654 (2022).

19. Paranjapye, A. et al. Autism spectrum disorder risk genes have convergent effects on transcription and neuronal firing patterns in primary neurons. Genome Res 35, 2433–2444 (2025).

20. Moore, S. M. et al. Setd5 haploinsufficiency alters neuronal network connectivity and leads to autistic-like behaviors in mice. Transl Psychiatry 9, 24 (2019).

21. Lukin, J. et al. Neuronal activity-dependent gene expression is stimulus-specific and changes with neuronal maturation. Front. Mol. Neurosci. 18, 1609772 (2025).

22. Lein, E. S. et al. Genome-wide atlas of gene expression in the adult mouse brain. Nature 445, 168–176 (2007).

23. Hubbard, K. S., Gut, I. M., Lyman, M. E. & McNutt, P. M. Longitudinal RNA sequencing of the deep transcriptome during neurogenesis of cortical glutamatergic neurons from murine ESCs. F1000Res 2, 35 (2013).

24. McCarthy, M. M. Estradiol and the Developing Brain. Physiological Reviews 88, 91–134 (2008).

25. Krebs-Kraft, D. L., Hill, M. N., Hillard, C. J. & McCarthy, M. M. Sex difference in cell proliferation in developing rat amygdala mediated by endocannabinoids has implications for social behavior. Proc. Natl. Acad. Sci. U.S.A. 107, 20535–20540 (2010).

26. McCarthy, M. M. & Nugent, B. M. Epigenetic Contributions to Hormonally-Mediated Sexual Differentiation of the Brain. J Neuroendocrinology 25, 1133–1140 (2013).

27. Nugent, B. M. et al. Brain feminization requires active repression of masculinization via DNA methylation. Nat Neurosci 18, 690–697 (2015).

28. Mulvey, B. et al. Molecular and Functional Sex Differences of Noradrenergic Neurons in the Mouse Locus Coeruleus. Cell Reports 23, 2225–2235 (2018).

29. Gegenhuber, B., Wu, M. V., Bronstein, R. & Tollkuhn, J. Gene regulation by gonadal hormone receptors underlies brain sex differences. Nature 606, 153–159 (2022).

30. Knoedler, J. R. et al. A functional cellular framework for sex and estrous cycle-dependent gene expression and behavior. Cell 185, 654–671.e22 (2022).

31. Fass, S. B. et al. Relationship between sex biases in gene expression and sex biases in autism and Alzheimer’s disease. Biol Sex Differ 15, 47 (2024).

32. Sun, E. D., Nagvekar, R., Pogson, A. N. & Brunet, A. Brain aging and rejuvenation at single-cell resolution. Neuron 113, 82–108 (2025).

33. Alpern, D. et al. BRB-seq: ultra-affordable high-throughput transcriptomics enabled by bulk RNA barcoding and sequencing. Genome Biol 20, 71 (2019).

34. Mitsuyoshi, S. et al. Expression of the proliferation-related Ki-67 mRNA in the early development of murine embryo. Biochem Biophys Res Commun 235, 191–196 (1997).

35. Ino, H. & Chiba, T. Expression of proliferating cell nuclear antigen (PCNA) in the adult and developing mouse nervous system. Molecular Brain Research 78, 163–174 (2000).

36. Graham, V., Khudyakov, J., Ellis, P. & Pevny, L. SOX2 Functions to Maintain Neural Progenitor Identity. Neuron 39, 749–765 (2003).

37. Vassilev, L. T. et al. Selective small-molecule inhibitor reveals critical mitotic functions of human CDK1. Proc. Natl. Acad. Sci. U.S.A. 103, 10660–10665 (2006).

38. Marangos, P. J., Schmechel, D., Zis, A. P. & Goodwin, F. K. The existence and neurobiological significance of neuronal and glial forms of the glycolytic enzyme enolase. Biol Psychiatry 14, 563–579 (1979).

39. Forrest, D. et al. Targeted disruption of NMDA receptor 1 gene abolishes NMDA response and results in neonatal death. Neuron 13, 325–338 (1994).

40. Kapitein, L. C. et al. NMDA Receptor Activation Suppresses Microtubule Growth and Spine Entry. J. Neurosci. 31, 8194–8209 (2011).

41. Greif, K. F., Asabere, N., Lutz, G. J. & Gallo, G. Synaptotagmin-1 promotes the formation of axonal filopodia and branches along the developing axons of forebrain neurons. Dev Neurobiol 73, 27–44 (2013).

42. Horton, A. C. et al. Polarized Secretory Trafficking Directs Cargo for Asymmetric Dendrite Growth and Morphogenesis. Neuron 48, 757–771 (2005).

43. Nakagawa, N. The neuronal Golgi in neural circuit formation and reorganization. Front. Neural Circuits 18, 1504422 (2024).

44. Lipton, J. O. & Sahin, M. The neurology of mTOR. Neuron 84, 275–291 (2014).

45. Panina, Y., Germond, A. & Watanabe, T. M. Analysis of the stability of 70 housekeeping genes during iPS reprogramming. Sci Rep 10, 21711 (2020).

46. Ho, K. H. & Patrizi, A. Assessment of common housekeeping genes as reference for gene expression studies using RT-qPCR in mouse choroid plexus. Sci Rep 11, 3278 (2021).

47. Hatakeyama, S. TRIM Family Proteins: Roles in Autophagy, Immunity, and Carcinogenesis. Trends in Biochemical Sciences 42, 297–311 (2017).

48. Malhotra, V., Serafini, T., Orci, L., Shepherd, J. C. & Rothman, J. E. Purification of a novel class of coated vesicles mediating biosynthetic protein transport through the Golgi stack. Cell 58, 329–336 (1989).

49. Son, G. & Han, J. Roles of mitochondria in neuronal development. BMB Rep 51, 549–556 (2018).

50. Iwata, R. et al. Mitochondria metabolism sets the species-specific tempo of neuronal development. Science 379, eabn4705 (2023).

51. Cao, J. et al. The single-cell transcriptional landscape of mammalian organogenesis. Nature 566, 496–502 (2019).

52. Fu, J. M. et al. Rare coding variation provides insight into the genetic architecture and phenotypic context of autism. Nat Genet 54, 1320–1331 (2022).

53. Hutton, S. R. & Pevny, L. H. SOX2 expression levels distinguish between neural progenitor populations of the developing dorsal telencephalon. Developmental Biology 352, 40–47 (2011).

54. Mercurio, S., Serra, L., Pagin, M. & Nicolis, S. K. Deconstructing Sox2 Function in Brain Development and Disease. Cells 11, 1604 (2022).

55. Kallur, T., Gisler, R., Lindvall, O. & Kokaia, Z. Pax6 promotes neurogenesis in human neural stem cells. Molecular and Cellular Neuroscience 38, 616–628 (2008).

56. Ypsilanti, A. R. & Rubenstein, J. L. R. Transcriptional and epigenetic mechanisms of early cortical development: An examination of how Pax6 coordinates cortical development. J Comp Neurol 524, 609–629 (2016).

57. Miyoshi, G., et al. *Prox1* Regulates the Subtype-Specific Development of Caudal Ganglionic Eminence-Derived GABAergic Cortical Interneurons. J. Neurosci. 35, 12869–12889 (2015).

58. Heuvelmans, A. M. et al. Modeling mTORopathy-related epilepsy in cultured murine hippocampal neurons using the multi-electrode array. Experimental Neurology 379, 114874 (2024).

59. Quintanilla, C. A. et al. High-density multielectrode arrays bring cellular resolution to neuronal activity and network analyses of corticospinal motor neurons. Sci Rep 15, 732 (2025).

60. Steinhoff, M. S., von Mentzer, B., Geppetti, P., Pothoulakis, C. & Bunnett, N. W. Tachykinins and their receptors: contributions to physiological control and the mechanisms of disease. Physiol Rev 94, 265–301 (2014).

61. Nässel, D. R., Zandawala, M., Kawada, T. & Satake, H. Tachykinins: Neuropeptides That Are Ancient, Diverse, Widespread and Functionally Pleiotropic. Front. Neurosci. 13, 1262 (2019).

62. Zimmer, A. et al. Hypoalgesia in mice with a targeted deletion of the tachykinin 1 gene. Proc. Natl. Acad. Sci. U.S.A. 95, 2630–2635 (1998).

63. Navratilova, E. & Porreca, F. Substance P and Inflammatory Pain: Getting It Wrong and Right Simultaneously. Neuron 101, 353–355 (2019).

64. Bartley, E. J. & Fillingim, R. B. Sex differences in pain: a brief review of clinical and experimental findings. Br J Anaesth 111, 52–58 (2013).

65. Failla, M. D. et al. Gender Differences in Pain Threshold, Unpleasantness, and Descending Pain Modulatory Activation Across the Adult Life Span: A Cross Sectional Study. The Journal of Pain 25, 1059–1069 (2024).

66. Tallent, M. K. et al. Cortistatin overexpression in transgenic mice produces deficits in synaptic plasticity and learning. Molecular and Cellular Neuroscience 30, 465–475 (2005).

67. Adams, J. M. et al. Somatostatin is essential for the sexual dimorphism of GH secretion, corticosteroid-binding globulin production, and corticosterone levels in mice. Endocrinology 156, 1052–1065 (2015).

68. Cordoba-Chacón, J. et al. Cortistatin Is a Key Factor Regulating the Sex-Dependent Response of the GH and Stress Axes to Fasting in Mice. Endocrinology 157, 2810–2823 (2016).

69. Jennings, K. J. & De Lecea, L. Neural and Hormonal Control of Sexual Behavior. Endocrinology 161, bqaa150 (2020).

70. Tisdale, R. K. et al. Biological sex influences sleep phenotype in mice experiencing spontaneous opioid withdrawal. Journal of Sleep Research 33, e14037 (2024).

71. Chaturvedi, S. M. et al. Chromosomal and gonadal sex have differing effects on social motivation in mice. Biol Sex Differ 16, 13 (2025).

72. Deegan, D. F. & Engel, N. Sexual Dimorphism in the Age of Genomics: How, When, Where. Front. Cell Dev. Biol. 7, 186 (2019).

73. Werner, R. J. et al. Sex chromosomes drive gene expression and regulatory dimorphisms in mouse embryonic stem cells. Biol Sex Differ 8, 28 (2017).

74. Syrett, C. M., Sierra, I., Berry, C. L., Beiting, D. & Anguera, M. C. Sex-Specific Gene Expression Differences Are Evident in Human Embryonic Stem Cells and During In Vitro Differentiation of Human Placental Progenitor Cells. Stem Cells Dev 27, 1360–1375 (2018).

75. Richardson, V., Engel, N. & Kulathinal, R. J. Comparative developmental genomics of sex-biased gene expression in early embryogenesis across mammals. Biol Sex Differ 14, 30 (2023).

76. Lu, T. & Mar, J. C. Investigating transcriptome-wide sex dimorphism by multi-level analysis of single-cell RNA sequencing data in ten mouse cell types. Biol Sex Differ 11, 61 (2020).

77. Deegan, D. F., Nigam, P. & Engel, N. Sexual Dimorphism of the Heart: Genetics, Epigenetics, and Development. Front. Cardiovasc. Med. 8, 668252 (2021).

78. McFarlane, L., Truong, V., Palmer, J. S. & Wilhelm, D. Novel PCR Assay for Determining the Genetic Sex of Mice. Sex Dev 7, 207–211 (2013).

79. Newman, A. M. et al. Determining cell type abundance and expression from bulk tissues with digital cytometry. Nat Biotechnol 37, 773–782 (2019).

80. Kolberg, L., Raudvere, U., Kuzmin, I., Vilo, J. & Peterson, H. gprofiler2 -- an R package for gene list functional enrichment analysis and namespace conversion toolset g:Profiler. F1000Res 9, ELIXIR-709 (2020).

81. Reimand, J., Kull, M., Peterson, H., Hansen, J. & Vilo, J. g:Profiler—a web-based toolset for functional profiling of gene lists from large-scale experiments. Nucleic Acids Research 35, W193–W200 (2007).

82. Koopmans, F. et al. SynGO: An Evidence-Based, Expert-Curated Knowledge Base for the Synapse. Neuron 103, 217–234.e4 (2019).

83. Bailey, T. L., Johnson, J., Grant, C. E. & Noble, W. S. The MEME Suite. Nucleic Acids Res 43, W39–W49 (2015).

84. Korotkevich, G. et al. Fast gene set enrichment analysis. Preprint at 10.1101/060012 (2016).

85. Bedogni, F. & Hevner, R. F. Cell-Type-Specific Gene Expression in Developing Mouse Neocortex: Intermediate Progenitors Implicated in Axon Development. Front. Mol. Neurosci. 14, 686034 (2021).

86. Ana Conesa, M. J. N. <Mj N. E. maSigPro. Bioconductor 10.18129/B9.BIOC.MASIGPRO (2017).

87. Cahill, K. M., Huo, Z., Tseng, G. C., Logan, R. W. & Seney, M. L. Improved identification of concordant and discordant gene expression signatures using an updated rank-rank hypergeometric overlap approach. Sci Rep 8, 9588 (2018).

